# Head Removal Enhances Planarian Electrotaxis

**DOI:** 10.1101/2022.01.10.475627

**Authors:** Ziad Sabry, Rui Wang, Aryo Jahromi, Christina Rabeler, William B. Kristan, Eva-Maria S. Collins

## Abstract

Sensing electric fields is an ability that certain animal species utilize for communication, hunting, and spatial orientation. Freshwater planarians move toward the cathode in a static electric field (cathodic electrotaxis). First described by Raymond Pearl more than a century ago, planarian electrotaxis has received little attention and the underlying mechanisms and evolutionary significance remain unknown. We developed an apparatus and scoring metrics for automated quantitative and mechanistic studies of planarian behavior upon exposure to a static electric field. Using this automated setup, we characterized electrotaxis in the planarian *Dugesia japonica* and found that this species responds to voltage instead of to current, in contrast to results from previous studies using other species. Because longer planarians exhibited more robust electrotaxis than shorter planarians, we hypothesized that signals from the head impede cathodic electrotaxis. To test this hypothesis, we took advantage of the regenerative abilities of planarians and compared electrotaxis in head and tail fragments of various lengths. We found that tail and trunk fragments electrotaxed while head fragments did not, regardless of size. However, we could restore cathodic electrotaxis in head fragments via decapitation, demonstrating that the presence of the head impaired cathodic electrotaxis. This result is in stark contrast to other stimulated behaviors such as phototaxis, thermotaxis or chemotaxis, which are weaker or absent in headless fragments. Thus, electrotaxis may be an important ability of headless planarian fragments to support survival prior to head regeneration.

**Summary statement:** We present a new method for quantitative studies of planarian electrotaxis and show that *Dugesia japonica* move toward the cathode. This behavior is enhanced by removal of the head.

## Introduction

Freshwater planarians are several mm long soft-bodied flatworms famous for their regenerative abilities (Rink, 2018). Planarians have a large repertoire of behaviors that can be used as readouts of brain function (Inoue *et al.*, 2015; Zhang *et al.*, 2019). These behaviors have been studied for over a century. Raymond Pearl (1903) was the first to write a comprehensive description of planarian behaviors, including ciliary driven gliding and musculature driven locomotion (peristalsis, scrunching), phototaxis, chemotaxis, and thermotaxis. Recently, planarians have experienced a resurgence as a neurobiology system because modern molecular biology techniques paired with computer vision now allow for mechanistic and quantitative studies of behavior. For example, it was shown that ciliary gliding depends on serotonergic signaling (Currie and Pearson, 2013), that peristalsis and scrunching are distinct gaits (Cochet-Escartin, Mickolajczk and Collins, 2015), with peristalsis resulting from non-functional cilia (Rompolas, Patel-King and King, 2010) and scrunching being a cilia-independent escape gait (Cochet-Escartin, Mickolajczk and Collins, 2015). Thermo-, photo-, and chemotaxis have been found to require the presence of a brain to sense their respective stimuli (Inoue *et al.*, 2015), whereas fission (Malinowski *et al.*, 2017; Goel *et al.*, 2021), scrunching (Cochet-Escartin, Mickolajczk and Collins, 2015), and avoidance of local near-UV stimulation (Le *et al.*, 2021) can occur without a brain.

Here, we characterize electrotaxis, another planarian behavior which was first described by Pearl over a century ago (Pearl, 1903) but, to the best of our knowledge, has not been rigorously revisited. Pearl showed that members of various planarian species *(Planaria maculata, dorotocephala, gonocephala;* **Table S1**) turn towards the negatively charged electrode (cathode) when an electrical field is applied (Pearl, 1903). He observed that the end of the planarian closest to the positively charged electrode (anode) contracted, comparable to the response observed by mechanical stimulation. Pearl interpreted this as evidence of the current acting directly on the muscles rather than interacting with sensory organs or cilia. Furthermore, he reported that planarians became “wholly or partially paralyzed in a very short time after the current begins to act, and as a consequence the reactions become feeble and indistinct” (Pearl, 1903). Unfortunately, no information on the duration of these experiments was provided, but this description of the planarians’ behavior suggests the use of strong electric fields. Finally, Pearl found that head pieces from transversely cut planarians behaved identically to intact worms while tail pieces displayed contraction on the anode-facing end but did not reorient or move towards the cathode.

Subsequent studies in the first half of the 20^th^ century by a handful of researchers on various planarian species (**Table S1**) confirmed that intact planarians either orient and move towards the cathode (Hyman and Bellamy, 1922; Robertson, 1927; Hyman, 1932) or assume a U- or W-shape, which allowed them to bring their head and, for certain species future heads at fission locations, closest to the cathode, depending on the strength of the electric field (Hyman and Bellamy, 1922; Hyman, 1932). For *Dugesia tigrina,* Pearl’s observation that the end of the planarian nearest the anode appears to contract was confirmed (Hyman and Bellamy, 1922; Robertson, 1927; Hyman, 1932). However, in contrast to Pearl’s findings, all planarian fragments (heads, trunks, tails) were reported to exhibit cathodic electrotaxis (Robertson, 1927; Fries, 1928; Marsh and Beams, 1952; Viaud, 1952a).

Different mechanisms have been proposed to explain planarian electrotaxis: 1) Direct action of electrical current on nerve or muscle cells (Pearl, 1903; Fries, 1928), 2) an intrinsic bioelectric gradient in the body of the animal, with a positively charged head and a negatively charged tail (Hyman and Bellamy, 1922; Robertson, 1927; Hyman, 1932), 3) a bioelectric gradient with a negatively charged head and a positively charged tail that causes electrophoresis of a negatively charged diffusible head inhibitor molecule with source at the head (Lange and Steele, 1978), or 4) directional differences in conductance (with less resistance in the head) and excitation along the anterior-posterior axis (Viaud and Medioni, 1951; Viaud, 1952b, 1952a). The existing experimental data, however, are insufficient to distinguish among these theories. Furthermore, because different researchers used different planarian species, varying experimental conditions and manual scoring metrics, which were rarely described in detail and may have suffered from experimenter bias (reviewed by (Jenkins, 1967) and summarized in **Table S1**), it is difficult to compile and interpret these previous findings. Therefore, we decided to revisit planarian electrotaxis with modern experimental tools and a quantitative and automated approach using the species *Dugesia japonica,* a popular species for planarian behavioral studies (Shomrat and Levin, 2013; Inoue, Yamashita and Agata, 2014; Inoue *et al.*, 2015; Sabry *et al.*, 2019; Zhang *et al.*, 2019; Ireland *et al.*, 2020; Le *et al.*, 2021).

## Materials and Methods

### Animal care

Asexual *Dugesia japonica* planarians were used for electrotaxis experiments. Planarians were kept in plastic containers filled with 0.5 g/L Instant-Ocean (IO) water (Spectrum Brands, Blacksburg, VA, USA) and stored at 18-20°C in temperature-controlled incubators (MIR-554, Panasonic, Kadoma, Osaka, Japan) in the dark when not used for experiments. Planarians were maintained following standard protocols (Dunkel, Talbot and Schötz, 2011), fed organic beef liver once a week, cleaned twice a week, and starved for at least one week before use in experiments.

### Electrotaxis arena setup

We developed an arena in which five planarians can be simultaneously imaged during exposure to a static electric field, with computer-controlled voltage strength and field direction. We designed a 60.0 mm long trough arena with an isosceles trapezoid cross section shape. A trapezoidal shape was chosen to ensure that planarians can be observed even when moving along the container boundaries, for which they have a preference (Akiyama, Agata and Inoue, 2015). The trough is 17.3 mm wide at the top, 4.4 mm wide at the bottom, and 10.0 mm in height **(Figure 1A)**. Five troughs (arenas) were milled into a transparent acrylic sheet, allowing up to five independent experiments to be run simultaneously. Electrodes that take up the cross section of the arena were cut out of a 3 mm thick aluminum sheet and adhered with cyanoacrylate glue to either end of each arena.

**Figure 1:**
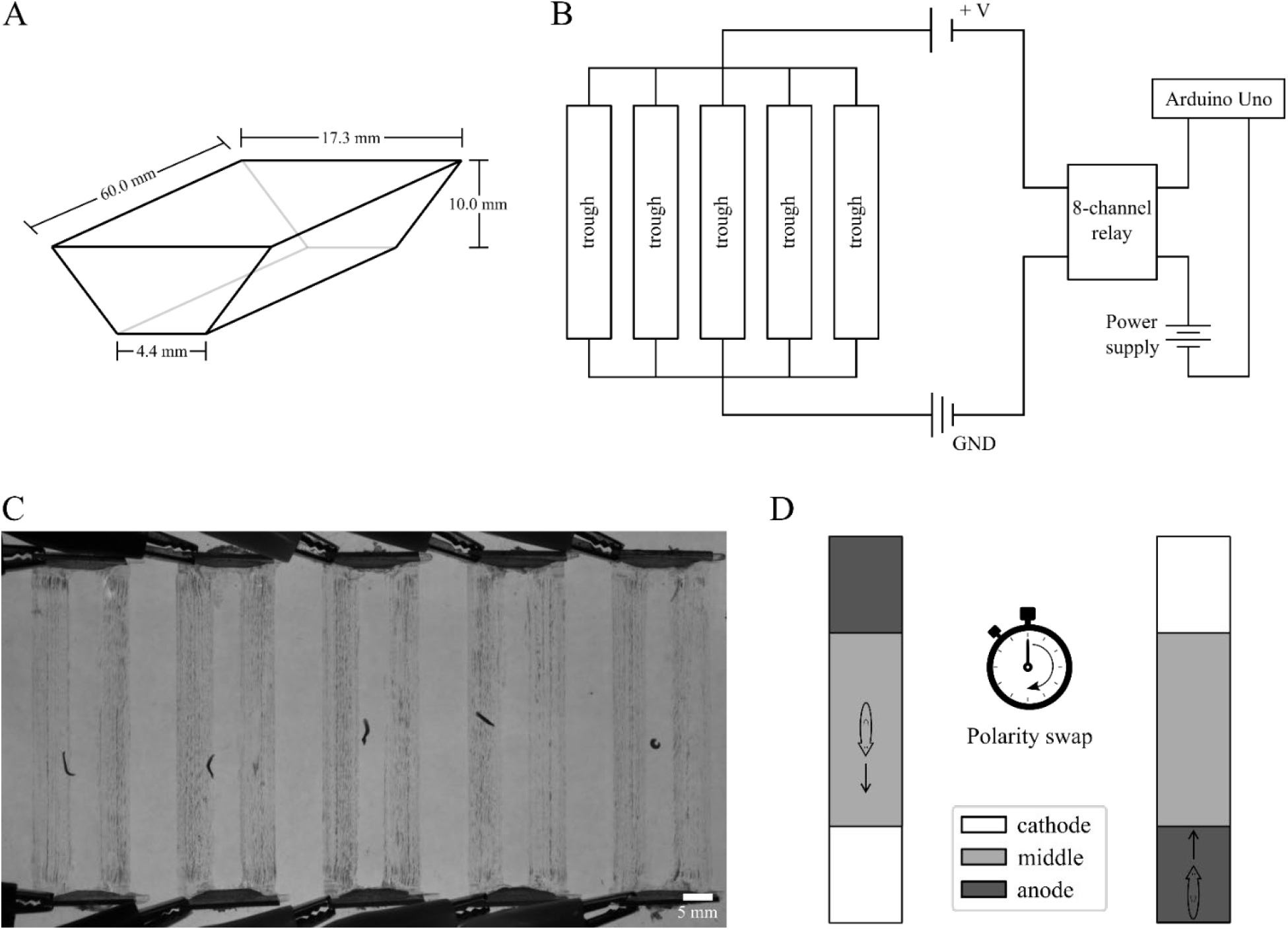
Schematics of planarian electrotaxis setup. (A) Schematic of one trough (arena) with an isosceles trapezoidal cross-section. (B) Circuit diagram of electrotaxis setup. (C) Representative image of planarians in the arenas backlit with a red electroluminescent panel. (D) Schematic showing a planarian in an arena. Planarians were dropped in the middle of each trough at the start of the experiment. The electrical polarity was reversed after half the experiment time had elapsed. White, grey, and dark-grey regions denote the cathode quadrant, middle two quadrants, and anode quadrant, respectively.

The five sets of electrodes were arranged in a parallel circuit configuration. An external 18 Volt DC 2.0 Linear Bench Power Supply (CircuitSpecialists, Tempe, AZ, USA) provided a voltage to the circuit. A voltage was supplied to each arena through an 8-Channel 5 V Relay Shield Module Board Optocoupler Module Arduino ARM PIC AVR (Jekewin (Amazon), Seattle, WA, USA) which was controlled by an Arduino Uno **(Figure 1B)**. The 8-channel relay was connected to the aluminum electrodes and to the power supply by wires with alligator clips. All other connections were made using wires on a half-size breadboard (Adafruit Industries, New York, NY, USA).

To record experiments, a Basler Ace acA640 camera (Basler AG, Ahrensburg, Germany) was mounted on a ring stand above the arenas. Images were recorded at a rate of 8 frames per second as JPEG image stacks. Circuit control via the Arduino and recording via the Basler camera were controlled through MATLAB (version R2019b, MathWorks, Natick, MA, USA). The arenas were backlit with a 20 cm x 15 cm red electroluminescent panel (Adafruit Industries, New York, NY, USA) to provide contrast between the planarian and background **(Figure 1C)**. All experiments were conducted with the room lights turned off. When filled with IO water with an applied voltage of 2 V, the arena would experience a voltage differential of 0.33 V/cm and an approximate current density of 0.077 mA/mm^2^.

### Experimental conditions

Each arena was evenly filled with 4 mL of IO water. After filling the arenas with water, a background image of the entire setup was taken to be used for later data processing. One planarian was carefully dropped into the middle of each of the five troughs using a Samco 691 transfer pipet (Thermo Fisher Scientific, Waltham, MA, USA). Once all planarians were approximately centered in their arenas, planarians were exposed to the electric field and recorded. After half the predetermined experiment time elapsed, the electrical polarity was swapped **(Figure 1D)**. Recording was terminated and the voltage was brought to 0 V at the conclusion of each experiment. Planarians were subsequently removed from the arenas and placed in a recovery container. Prior to the beginning of another experiment, IO water was drained from each arena and arenas were wiped down with a paper towel to remove any mucus trails. All experiments were conducted at room temperature. For the voltage sweep, planarians were released approximately in the middle of their troughs at experiment onset (**Figure 1**). The experiment was 120 seconds in duration, with a polarity swap at 60 seconds (3 technical replicates with 5 planarians each in individual troughs for each voltage). To determine whether planarians responded to electrical current or to voltage, troughs were filled with 4 mL of either IO or IO or ultrapure (Milli-Q; MQ) water. Experiments were run at 4 V for 90 seconds without a polarity swap. The higher voltage of 4 V was chosen to achieve: 1) a high voltage, high current condition (using IO water) and 2) a high voltage, low current condition (using ultra-pure water).

### Temperature, convective currents, and pH tests

Temperature and pH differentials were measured when a 2 V electric field was applied to the arena, filled with 4 mL of IO water, for 360 seconds, with a polarity swap at 180 seconds (**SI Figure 1**). The pH was measured to be approximately 6.5 both before and after the electric field was applied using pH test strips (Whatman, Maidstone, United Kingdom). To measure temperature, an image of the arenas was taken before and after the electric field was applied using a FLIR infrared camera (FLIR Systems, Wilsonville, OR USA). To test for convection, a drop of food coloring dye (Gel Spice Company Inc., Bayonne, NJ USA) was placed at the initial anode before a 2 V electric field was applied to an arena for 180 seconds, with a polarity swap at 90 seconds. As comparison, a drop of food coloring dye was placed in the same region of a different arena with a 0 V electric field. An image of the arenas was taken before and after the electric field was applied.

### Amputation experiments

For all experiments involving amputations, transverse amputations were used. To generate head and tail pieces, planarians were amputated either just above (pre-pharyngeal) or just below (post-pharyngeal) the pharynx using a sterile razor blade. For experiments involving trunk pieces, planarians were amputated both pre-pharyngeally and just below the auricles. For successive amputations, cuts transversally to the head-tail axis were administered in series. After each amputation, planarians were given at least one day to heal prior to conducting electrotaxis experiments. Because small tail pieces are less mobile than intact worms (Inoue, Yamashita and Agata, 2014; Inoue *et al.*, 2015), we increased the duration of the experiment when assessing post-pharyngeally cut tails. Head pieces and pre-pharyngeal tails were exposed to the electric field for 240 s, with an electrical polarity swap at 120 s; post-pharyngeal tails were exposed for 360 s with a polarity swap at 180 s.

### Raw image data processing

Raw image data was imported into Fiji (Schindelin *et al.*, 2012) for background subtraction. The 5 arenas were separated into 5 image stacks using the rectangle tool to draw equal-sized rectangles around each arena and duplicating into individual image stacks. The arena background was subtracted from each of the five image stacks using the background image taken at the start of each experiment. The 5 stacks were then saved separately.

### Data analysis and statistics

Processed frames were imported into MATLAB and binarized by manually setting a threshold that encompassed only the planarian. The center of mass of each planarian was then tracked in each frame using custom MATLAB code as previously described (Talbot and Schötz, 2011). The time spent in the arena quadrants and the fraction of time spent moving towards the cathodes for each planarian were outputted and compiled into a spreadsheet. To quantify the planarian’s response to the electrical field, we calculated the fraction of time spent in the quadrant containing the cathode during the first half of the experiment *f_cat-1_,* time spent in cathode quadrant divided by total time with cathode at that location) and the fraction of time spent moving toward the cathode (*f_mov-1_*, time spent moving towards cathode divided by total time with cathode at that location). After the polarity swap, the same two parameters were calculated again *(f_cat-2_, f_mov-2_).* The fraction of experiment time spent in the middle two quadrants and in the anode quadrant (before and after the electrical polarity swap) were also recorded.

Responses to the electrical field were tested using analysis of variance (ANOVA) models. Response variables were proportions, either of trial time spent in the cathode zone or of trial time spent moving toward the cathode, before or after polarity swaps. Differences in these values between controls measured without a voltage applied and treatment group worms with a voltage applied constituted electrotaxis. For the initial test of electrotaxis at varying voltage levels, a oneway ANOVA was used with voltage as a predictor variable. Significant effects of voltage were followed up with Dunnett’s post-hoc tests of 0 V controls against the non-zero voltages. Experiments with additional treatments were analyzed with factorial ANOVA. Since the difference between 0 V and 2 V was the indication of electrotaxis, the effect of other predictors (such as worm size) on electrotaxis was indicated by a significant interaction between voltage and the other predictors (that is, the amount of electrotaxis depends on worm size if the voltage x size interaction is significant). When significant interactions were detected the difference between 0 V and 2 V groups at the levels of the other predictor were used as post-hoc procedures, using Tukey’s method to account for multiple comparisons. Post-hoc comparisons were only conducted following a significant ANOVA, so p-values for post-hoc comparisons of group means will be reported with the post-hoc procedure identified.

The successive cuts experiment used the same planarians with treatments that changed each day. These repeated-measures data were analyzed using the successive treatments as a within-subjects factor. The within-subjects treatments differed between two groups of worms, so the group was used as a between-subjects factor. The interaction of the within and between-subjects factors indicated that the two groups differed in their responses to the two different treatment sequences. The same worms were tested at 0 V and 2 V to test for electrotaxis on each day, so post-hoc comparisons between the voltage levels were done with paired t-tests, and significance was assessed with a Bonferroni-adjusted alpha level (six comparisons were made, so p-values needed to be less than 0.05/6 = 0.008 to be considered statistically significant for the post-hoc paired t-tests).

Proportions often violate assumptions of normality and homoscedasticity, so these assumptions were tested prior to each analysis. When one or more assumptions were violated, we used non-parametric randomization tests to confirm that statistical significance of model terms was not affected. If significance was unchanged by using randomization tests, then post hoc procedures were conducted as usual, using either Tukey or Dunnett tests. All analysis was done with the R statistical computing language (version 4.1.2, R Core Team) and extension libraries. Post-hoc procedures were done with library emmeans (version 1.7.0). Randomization tests were done using library lmPerm (2.1.0). Homoscedasticity was tested with a Breusch-Pagan test from the lmtest library (0.9-38). Repeated measures analysis was done with the car library (3.0-11). The data, analysis in R and respective figures can be downloaded from the Collinslab github (https://github.com/Collinslab-swat/Planarian-electrotaxis).

## Results

### D. japonica exhibits cathodic electrotaxis at 2 V without overt adverse effects

To determine what field strength was necessary to induce electrotaxis, we conducted a voltage sweep **(Table 1)**. At 0 V, planarians moved randomly (**SI Figure 2**) and one would expect them to spend 25% of the experimental time in each quadrant. We found that they spent approximately 1/4-1/3 of the experimental time in the quadrants containing the cathode **(Table 1)**. The increased time spent in the quadrants containing the electrodes compared to the two middle quadrants likely results from planarians exhibiting wall preference (Akiyama, Agata and Inoue, 2015), as the electrode containing quadrants have more walls than the middle two quadrants (**Figure 1C, D)**. Planarians did not exhibit a preference for movement toward either electrode at 0 V; they moved toward and away from the cathode before and after polarity swap at equal rates **(Table 1; Movie 1)**. When an electric field of 1 V was applied, planarians spent more time moving toward and staying in the cathode containing quadrants, but the increase was not significant compared to 0 V (Dunnett’s test, p = 0.587). At 1.5-2 V, planarians reoriented themselves and moved toward the cathode (**Movie 1** shows planarian behavior at 2 V) and spent > 50% of the experimental time in the cathode-containing quadrants **(Table 1,** Dunnett’s test, p < 0.001). Planarians did not spend significantly more time moving toward the first cathode at 1.5 V but did so at 2 V and higher voltages **(Table 1,** Dunnett’s test, p = 0.16 for 1.5 V, p = 0.006 for 2 V,p < 0.001 for 3 V and 4 V). Once the polarity was swapped, planarians had a longer distance to move toward the new cathode because they predominantly started in the most distant quadrant, at cathode 1 **(Figure 1D)**. We observed that following the polarity swap, time moving towards cathode 2 *(f_mov-2_)* appears to be a more consistent and sensitive measure for all voltages >1.5 V than time spent at the cathode *(f_cat-2_*), **Table 1**). This may be because planarians require longer to arrive at the second cathode from the most distant quadrant, causing time spent at the second cathode to be artificially low, whereas the time spent moving toward the cathode is relatively unchanged after the polarity swap.

**Table 1.** Parameters for baseline experiments. For each voltage tested, N=15 planarians (6.4 mm - 11.1 mm in length) were used in 3 experimental replicates with N=5. Voltage values are reported to ± 0.01 V; current values are averages of 4 measurements. Electrotaxis parameter values are reported as mean ± standard error. * denotes p < 0.05, ** denotes p < 0.01 and *** denotes p < 0.001 differences from the 0 V control using Dunnett’s post-hoc comparisons.

| Voltage | Current | $f_{cat-1}$ | $f_{mov-1}$ | $f_{cat-2}$ | $f_{mov-2}$ |
| --- | --- | --- | --- | --- | --- |
| 0 V | 0 A | $0.22 \pm 0.06$ | $0.49 \pm 0.03$ | $0.36 \pm 0.04$ | $0.51 \pm 0.03$ |
| 1 V | 0.03 mA | $0.33 \pm 0.06$ | $0.53 \pm 0.02$ | $0.39 \pm 0.04$ | $0.55 \pm 0.02$ |
| 1.5 V | 4.4 mA | $0.64 \pm 0.07^{***}$ | $0.56 \pm 0.02$ | $0.56 \pm 0.03^{**}$ | $0.63 \pm 0.02^{**}$ |
| 2 V | 8.4 mA | $0.65 \pm 0.07^{***}$ | $0.63 \pm 0.02^{***}$ | $0.54 \pm 0.05^*$ | $0.72 \pm 0.01^{***}$ |
| 3 V | 16.6 mA | $0.80 \pm 0.04^{***}$ | $0.76 \pm 0.03^{***}$ | $0.10 \pm 0.05^{***}$ | $0.72 \pm 0.02^{***}$ |
| 4 V | 25.1 mA | $0.72 \pm 0.06^{***}$ | $0.79 \pm 0.03^{***}$ | $0.08 \pm 0.03^{***}$ | $0.88 \pm 0.02^{***}$ |

While planarians exhibited electrotaxis at 3-4 V, they also exhibited vigorous head turning and oscillatory behavior (**Movie 2** shows planarian behavior at 4 V). These behaviors caused the planarians to move more slowly toward the cathode. They still were able to reach the first cathode because they only had to traverse half of the trough but failed to reach the cathode after the polarity switch because they needed to travel the whole distance and the adverse effects increased over time. Because of these adverse effects at higher voltages, we conducted all further experiments at 2 V.

Next, we investigated whether the planarians sensed the electric field directly or they reacted to secondary effects induced by the field, such as gradients in pH, temperature, and convective currents, which can affect planarian behavior (Inoue, Yamashita and Agata, 2014; Ross *et al.*, 2018; Sabry *et al.*, 2020). We did not find significant effects of any of these factors **(SI Figure 1)**. Thus, planarian movement toward the cathode is a direct response to the electric field.

To determine whether planarians responded to electrical current or to voltage, we tested the planarians’ response to 4 V in either IO or ultrapure (MQ) water. Since the *f_cat-2_* parameter is not indicative of electrotaxis ability at 4 V as seen in **Table 1**, we calculated electrotaxis parameters without an electrical polarity swap. At 4 V, the current across a single trough of IO and ultra-pure water were measured to be 25.1 mA and 3.3 μA, respectively (averaged over 4 measurements). In IO water at 4 V, planarians spent significantly more time in the quadrant containing the cathode **(SI Figure 1C)** as well as spent more time moving towards the cathode compared to 0 V **(SI Figure 1D)**. The same trend was observed when planarians were placed in MQ water **(SI Figure 1C-D)**, demonstrating that planarian movement toward the cathode is not due to the electrical current (which differed by 4 orders of magnitude) but due to voltage. We will refer to this behavior as cathodic electrotaxis in subsequent sections.

### Planarian body length is predictive of cathodic electrotaxis ability

It has been previously shown that planarian behaviors such as locomotor velocity can be size dependent (Talbot and Schötz, 2011). To determine whether size also plays a role in planarian cathodic electrotaxis, experiments were run at 2 V on N=94 planarians that ranged in size from 2.0 - 12.4 mm. Planarians were classified as either “small”, “medium”, or “large”**(SI Figure 2A**). Planarians in the large size class (7.6-12.4 mm) spent significantly more time at 2 V than at 0 V moving toward and staying in the quadrant containing the cathode, before and after the electrical polarity swap **(Figure 2A, B, SI Figure 2B,** Tukey post-hocs, p < 0.001). Planarians in the medium size class (4.6-7.5 mm) spent significantly more time at 2 V than at 0 V moving toward and staying near the first cathode **(Figure 2C, D,** Tukey post-hocs, p < 0.003). After polarity reversal, medium sized planarians spent significantly more time moving towards the second cathode when voltage was applied (Tukey post-hocs, p < 0.001), although the time spent in the quadrant containing the second cathode was not significantly different between 0 V and 2 V **(Figure 2C**, Tukey post-hocs, p = 0.077).

**Figure 2:**
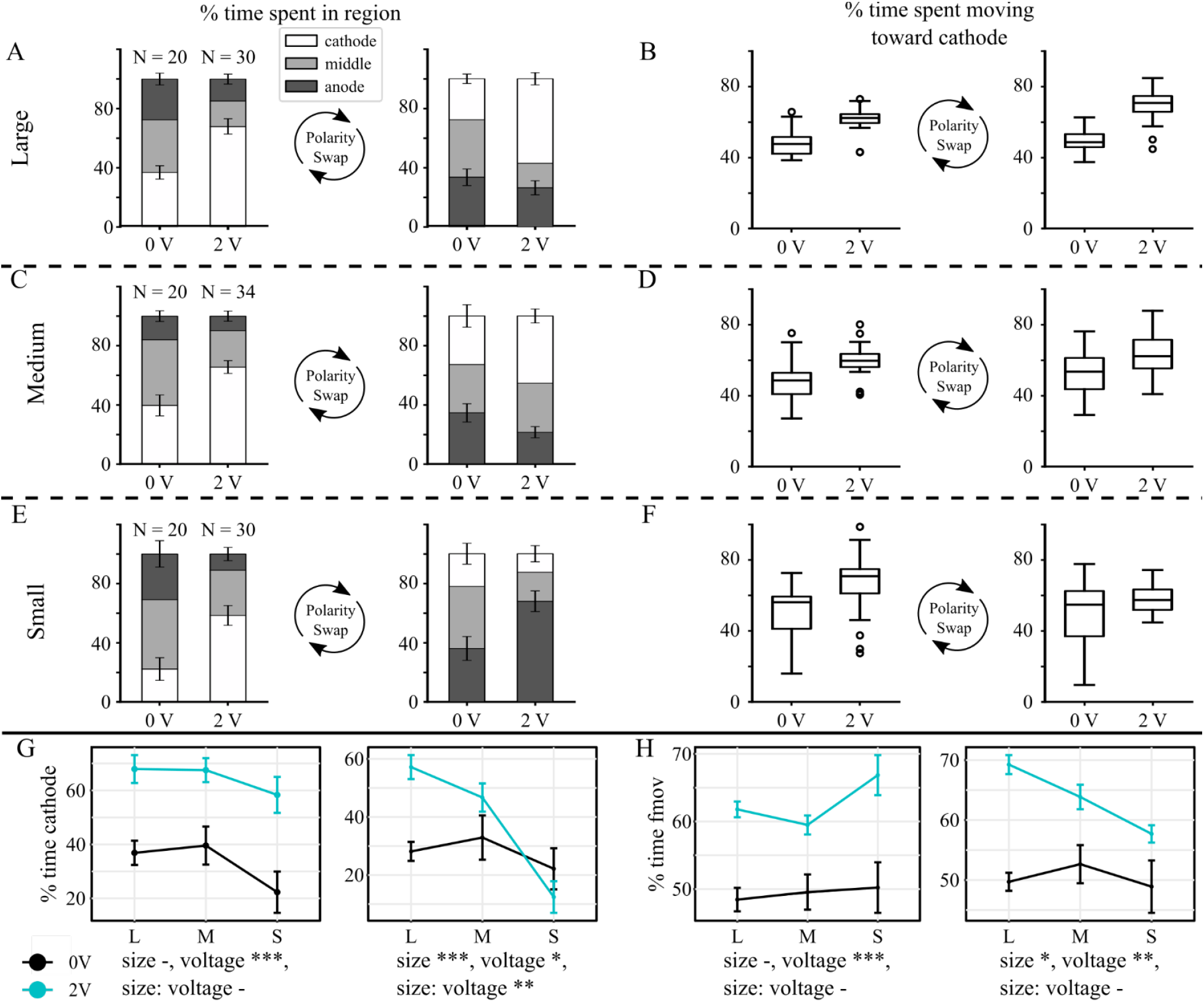
Size and electrotaxis ability. (A, C, E) Segmented bar plots showing the percent experiment time, before and after the electrical polarity swap, spent in the cathode quadrant, anode quadrant, and middle two quadrants for (A) large, (C) medium, and (E) small sized planarians. Error bars denote standard error. (B, D, F) Box-and-whisker plots showing the percentage of experiment time, before and after the electrical polarity swap, spent moving toward the cathode for (B) large, (D) medium, and (F) small sized planarians. Open circles denote outliers. G. Interaction plots showing % time spent at the cathode for small, medium, and large planarians at 0 V and 2 V, before and after electrical polarity swap. H. Interaction plots showing % time moving towards the cathode for small, medium, and large planarians at 0 V and 2 V, before and after electrical polarity swap. *** denotes p < 0.001, ** denotes p < 0.01, * denotes p < 0.05, - denotes p > 0.05. Shown are mean values and error bars denote standard error.

Small planarians (2.0-3.5 mm) spent significantly more time moving towards and staying near the first cathode at 2 V than at 0 V **(Figure 2E, F,** Tukey post-hocs, p < 0.001). After the electrical polarity swap, small planarians spent significantly more time moving towards the second cathode (Tukey post-hocs, p = 0.011) but there was no difference in time spent in the quadrant containing the second cathode **(Figure 2E, F**, Tukey post-hocs, p = 0.228). This difference in behavior for smaller versus larger planarians was not due to differences in motility. While smaller planarians are known to move slower than larger planarians (Hagstrom *et al.*, 2015) and we observed differences in speed ((1.70 +/- 0.05) mm/s for large planarians and (1.02 +/- 0.04) mm/s for small planarians (mean +/- std); N= 30 and N=25, respectively), there was sufficient time (90 sec) for small planarians to travel the length of the trough (60 mm). However, in contrast to large planarians, small planarians did not move toward the new cathode after the polarity swap but wandered around the anode containing quadrant (**Figure 2E** and **SI Figure 2**).

### Cathodic electrotaxis is a brain-independent behavior

To test whether cathodic electrotaxis requires key anatomical structures, such as the head, tail, auricles, and pharynx, we bisected planarians into head and tail pieces either anterior to (pre) or posterior to (post-) the pharynx **(Figure 3 A,B)**. Amputated planarians were allowed one day to heal before assaying for electrotaxis ability. Because size and motility affect the time spent at the cathodes and because the behavior at the cathodes is also influenced by other factors, such the planarians’ wall preference behavior (Akiyama, Agata and Inoue, 2015), we focused on the motility parameters *f_mov-1,2_* for all further analyses.

**Figure 3:**
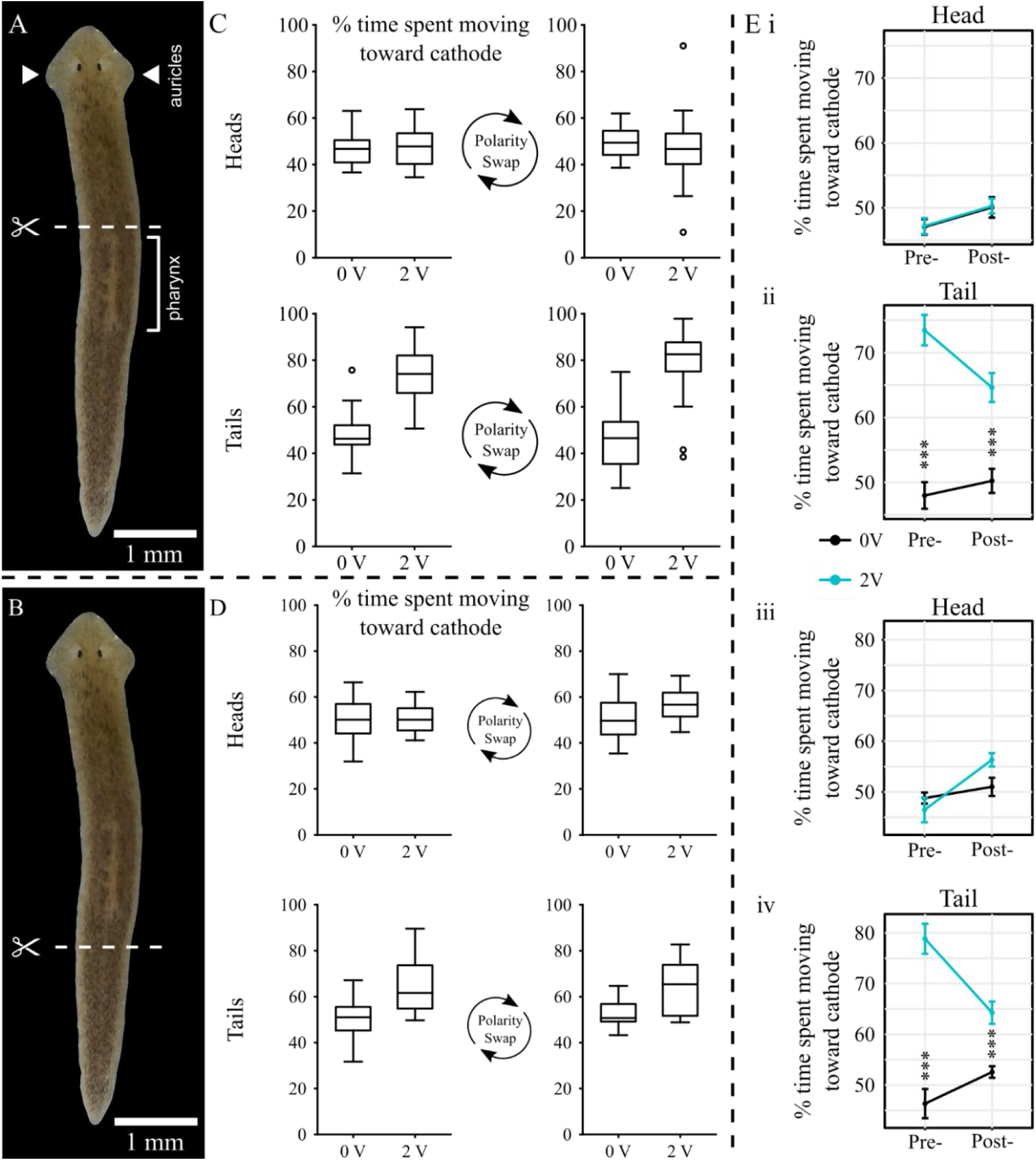
Electrotaxis behavior of head pieces depends on cut location. (A) Schematic showing the site of pre- pharyngeal amputations. Pharynx location indicated by bracket; auricles indicated by white arrows. (B) Schematic showing the post-pharyngeal cut location. (C) Box-and-whisker plots showing the percentage of experiment time, before and after the electrical polarity swap, spent moving toward the cathode. Open circles denote outliers. (D) Percent experiment time spent moving toward the cathode for post-pharyngeally cut (i) head and (ii) tail pieces. (E) Comparisons of time spent moving towards the cathode for pre- and post-pharyngeal head and tail fragments. i, ii. Before polarity swap, iii, iv. After polarity swap. * denotes p < 0.05, ** denotes p < 0.01, and *** denotes p < 0.001 from the respective 0 V controls as calculated using Tukey post-hoc comparisons. Shown are mean values and error bars denote standard error.

If electrotaxis ability was solely size dependent, we would expect to find that larger fragments electrotax more robustly than smaller ones, independent of their head or tail identity. However, we found that tail pieces retained the ability to electrotax independent of size (Tukey post-hocs, p < 0.001; **SI Figure 3**), whereas head pieces did not exhibit cathodic electrotaxis regardless of amputation location **(Figure 3 C-E**, Tukey post-hocs, p = 0.378 for pre-pharyngeally amputated pieces, p = 0.063 for post-pharyngeally amputated pieces; **SI Figure 3)**. As planarian tail pieces lack brains yet still maintain the ability to electrotax, these results demonstrate that the planarian brain or the auricles are not required for electrotaxis. Furthermore, because tail pieces with and without the pharynx electrotaxed (**SI Figure 4**), these experiments show that the pharynx is not required for cathodic electrotaxis. Moreover, small post-pharyngeal tails electrotaxed more robustly than larger pre-pharyngeal heads (**Figure 3E**), suggesting that the presence of a head may impair electrotaxis ability.

We tested the hypothesis that the relative size of the head to the body may affect electrotaxis by taking head and body length measurements from our data (**Figure 4A).** In support of this idea, we observed a weak interaction between head proportion and time spent moving towards the second cathode (*f_mov-2_*) (**Figure 4B**). Therefore, we decided to use a direct experimental approach to dissect the role of the head for the electrotaxis ability of individual planarians.

**Figure 4.**
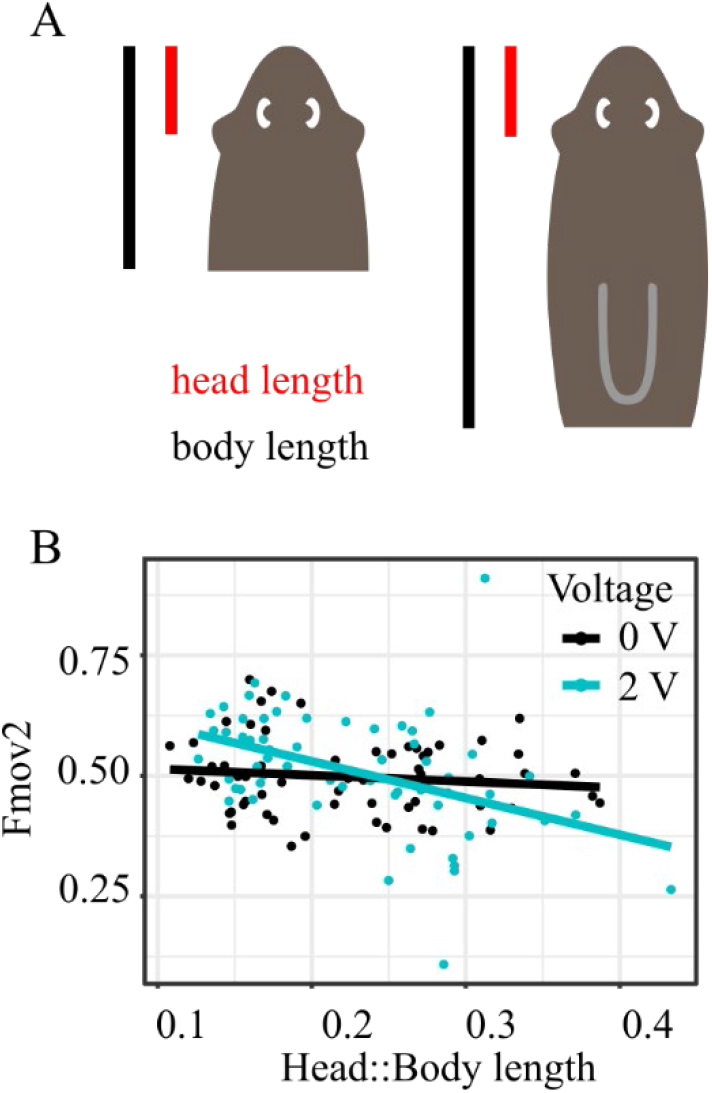
Head proportion does not have a major impact on electrotaxis behavior. A. Schematic showing head fragments and calculation of head proportion. B. Interaction plots between head length to body length ratio and time spent moving towards second cathode.

### Head removal restores electrotaxis behavior in planarian fragments

First, we quantified electrotaxis in pre-pharyngeal heads and trunks **(SI Figure 5)**. These animals were exposed to a 2 V electric field for 240 s with a polarity swap at 120 s. We then decapitated the heads, removed an equivalent tissue fragment from the anterior end of the trunks, and re-evaluated electrotaxis after 24 h, to allow for healing. We found that pre-pharyngeal heads do not electrotax, but this ability is restored by decapitation **(SI Figure 5).**Because one could argue that (a) the planarians may have differed in their ability to electrotax from the beginning and (b) that any anterior cut may restore electrotaxis, we repeated this experiment using successive cuts on large planarians with tracking of individual animals that we verified to have electrotaxis ability. Planarians were cut post-pharyngeally, allowed to heal for 24 h, and split into two groups (**Figure 5A**). Subsequently group 1 had a small amount of tissue removed from the posterior end and group 2 was cut pre-pharyngeally. On day 2, both groups were again assessed for electrotaxis, after which group 1 was cut pre-pharyngeally and had a small amount of tissue removed from the tip of the nose while group 2 was decapitated. On day 3, both groups were assessed for a final time (**Figure 5A**). Members of group 1 retained a head throughout the experiment, whereas members of group 2 lost their head in the third amputation. Each group received the same number of posterior and anterior cuts, to account for any changes that may be introduced by amputation.

**Figure 5:**
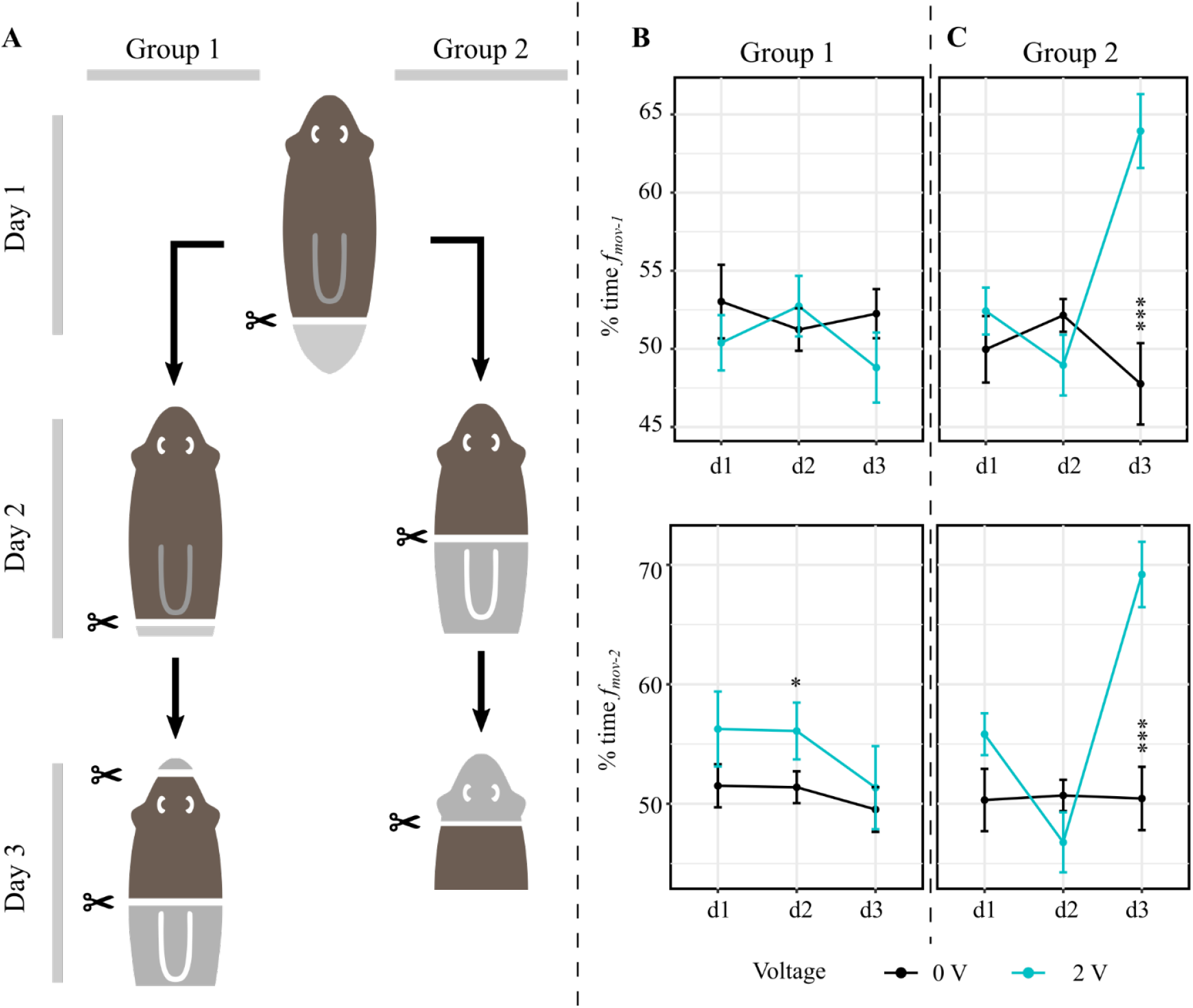
Electrotaxis metrics are dependent on presence or absence of a head. A. Schematic of experimental procedure. Notably, the indicated cuts were performed the day prior to the experimental day indicated in the text. B. Time spent moving towards the cathode on each day, Group 1. C. Time spent moving towards the cathode on each day, Group 2. * denotes p < 0.05, ** denotes p < 0.01, and *** denotes p < 0.001 from the respective 0 V controls as determined by post hoc comparisons. Shown are mean values and error bars denote standard error.

We found that post-pharyngeally cut heads (day 1) showed no electrotaxis, consistent with earlier experiments. Within Group 1, post-pharyngeal heads with an additional posterior wound (day 2) showed weak electrotaxis. Pre-pharyngeal amputation coupled with an anterior wound (day 3) caused loss of electrotaxis **(Figure 5B)**. Similarly, within group 2, pre-pharyngeal heads (day 2) failed to show electrotaxis. However, trunk pieces formed by subsequent decapitation (day 3) showed clear recovery of electrotaxis (**Movie 3**) with statistically significant changes in both *f_mov-1_* and *f_mov-2_* **(Figure 5C)**. A non-decapitating anterior wound (Group 1) did not restore electrotaxis. Based on these results and length measurements (**SI Figure 3**), we conclude that it is not fragment/planarian size but the presence or absence of a head that is the strongest factor determining planarian cathodic electrotaxis.

## Discussion

Using an automated experimental setup free of experimenter bias or other external influences, our data show that planarians respond to the electric field rather than to other environmental cues. This is an important distinction to make as planarians are known to sense temperature and chemical gradients (Inoue, Yamashita and Agata, 2014; Inoue *et al.*, 2015) and electric fields in water can generate thermal, pH, and convective effects (Gunji and Washizu, 2005; May and Hillier, 2005). While these effects should be minimal given the small voltage applied, they were important to test as it is unclear how sensitive planarians are. The voltages in previous studies (Pearl, 1903; Hyman and Bellamy, 1922; Robertson, 1927; Fries, 1928; Hyman, 1932), whether directly reported or estimated based on current and arena dimensions (**SI Table 1**), are much larger than the 2 V we used in our experiments.

The high and variable voltages involved likely account for some of the variability in the reported results, as well as the observed dramatic behaviors such as planarians curling up on their sides (Hyman and Bellamy, 1922; Hyman, 1932), scrunching (Robertson, 1927), paralysis and death (Pearl, 1903). These prior studies largely assumed that the response was elicited by current (Pearl, 1903; Hyman and Bellamy, 1922; Robertson, 1927; Fries, 1928; Hyman, 1932) (**SI Table 1**). The finding that planarians electrotax similarly in both ionized and deionized water despite a current difference of 4 orders of magnitude demonstrates that *D. japonica* planarians sense and respond to voltage and not to electrical current. The distinction between voltage and current informed our subsequent experimental design and is key to future efforts to determine the mechanism underlying planarian electrotaxis.

We showed that electrotaxis is not tied to a specific anatomical structure via amputation experiments. We assayed the role of the central nervous system (comprised of brain and nerve cords), and sensory structures such as the auricles and pharynx, which are used in chemotaxis (Asano *et al.*, 1998; Miyamoto *et al.*, 2020). Pre-pharyngeally cut tail pieces lack the brain and auricles, while post-pharyngeally cut tail pieces also lack the pharynx. The presence of electrotactic behavior in both types of tail pieces shows that the brain, pharynx, and auricles are not required. Furthermore, the presence of the ventral nerve cords seems insufficient to cause cathodic electrotaxis, because head fragments that also contain ventral nerve cords did not move to the cathode. Thus, voltage sensing, and subsequent directed motion cannot be attributed to specific anatomical structures but rather depends on a broadly distributed or graded property throughout the body. In addition, our voltage sweep showed that cathodic electrotaxis does not result from direct electrical action on either the cilia or muscles, because we were able to elicit the behavior in planarians gliding (1.5-2 V) and using musculature-driven locomotion (scrunching) (3-4 V).

Our observation that electrotaxis is weaker in smaller worms is interesting, as smaller planarians do not represent a different life stage where certain structures or tissues might be absent or immature. However, our results from varying sizes of both intact planarians and fragments show that electrotaxis ability is not a consequence of size (**SI Figure 3**). Instead, we find that a fragment containing a head lacks the ability to electrotax whereas a similarly sized fragment without a head retains this ability. This finding is the opposite of what was reported in the literature for other planarian species (**Table SI 1**), wherein it was found that head pieces in an electric field behaved more like intact planarians than other fragments (Pearl, 1903; Fries, 1928).

Behavioral differences among species are known to exist for other stimulated behaviors (Ireland *et al.*, 2020) and it is possible that the electrotaxis response of *D. japonica* differs from the other planarian species previously studied. An alternative explanation is that our use of lower field strength to avoid the adverse effects of electric field exposure described in the literature (scrunching, curling, paralysis, death (Pearl, 1903; Hyman and Bellamy, 1922; Robertson, 1927; Hyman, 1932)) elicited more differentiated behaviors.

Previous work on planarian electrotaxis has attributed the worms’ reaction to electric fields to direct action of the electric current on the muscles (Pearl, 1903; Fries, 1928), intrinsic bioelectric gradients of the animal (Robertson, 1927; Hyman, 1932; Lange and Steele, 1978) or head-to-tail differences in electrical conductance (Viaud and Medioni, 1951; Viaud, 1952a). Lange & Steele proposed an electrochemical model for axial patterning and measured the intrinsic bioelectric gradient. They reported that the head was negatively charged relative to the body, with a posteriorly increasing positive charge toward the tail (Lange and Steele, 1978). Based on these data, they proposed that the head-to-tail bioelectric gradient caused electrophoresis of a negatively charged head inhibitor molecule that is produced in the brain. Thus, according to their model, a static bioelectric gradient exists that is superimposed by a dynamic concentration gradient of a negatively charged morphogen that travels head to tail. Upon decapitation, a piece whose bioelectric gradient was aligned with an external electric field would thus experience a positive anterior relative to its posterior and migrate toward the cathode, as observed in the classical patterning experiments by Marsh & Beams (Marsh and Beams, 1952).

Conversely, Hyman proposed that the bioelectric gradient correlates with a metabolic gradient and because the head was more metabolically active, the head region was positively charged compared to the body (Hyman, 1932). Recent work (Durant *et al.*, 2017) using the DiBAC4(3) voltage reporter (Oviedo *et al.*, 2008) showed that the very tip of the head region is relatively depolarized (positively charged). While this result seems to support Hyman’s model, it does not directly contradict the measurements of Steele & Lange, given the coarse nature of their measurements and the observation that most of the head does not appear depolarized in the DiBAC experiments. DiBAC experiments also showed that trunk pieces have polarity with anterior wounds being more positively charged than posterior wounds (Durant *et al.*, 2019), in agreement with both model predictions and the observed cathodic electrotaxis (Marsh and Beams, 1952). A fourth explanation for electrotaxis was provided by Viaud & Medioni who reported that electrical conductance and excitation was consistently greater and the threshold for a response to current was lower when the planarian’s head was facing the cathode than when it faced the anode (Viaud and Medioni, 1951). This observation was reproduced in head and tail fragments (Viaud, 1952a).

How do these different explanations perform in the light of our experimental data? We can rule out the direct action of current on the musculature as the driving force for electrotaxis because we were able to elicit electrotaxis at 2 V without musculature driven locomotion. The models that propose anterior-posterior bioelectric or conductance gradients similarly cannot explain all the data (**Figure 6**).

**Figure 6:**
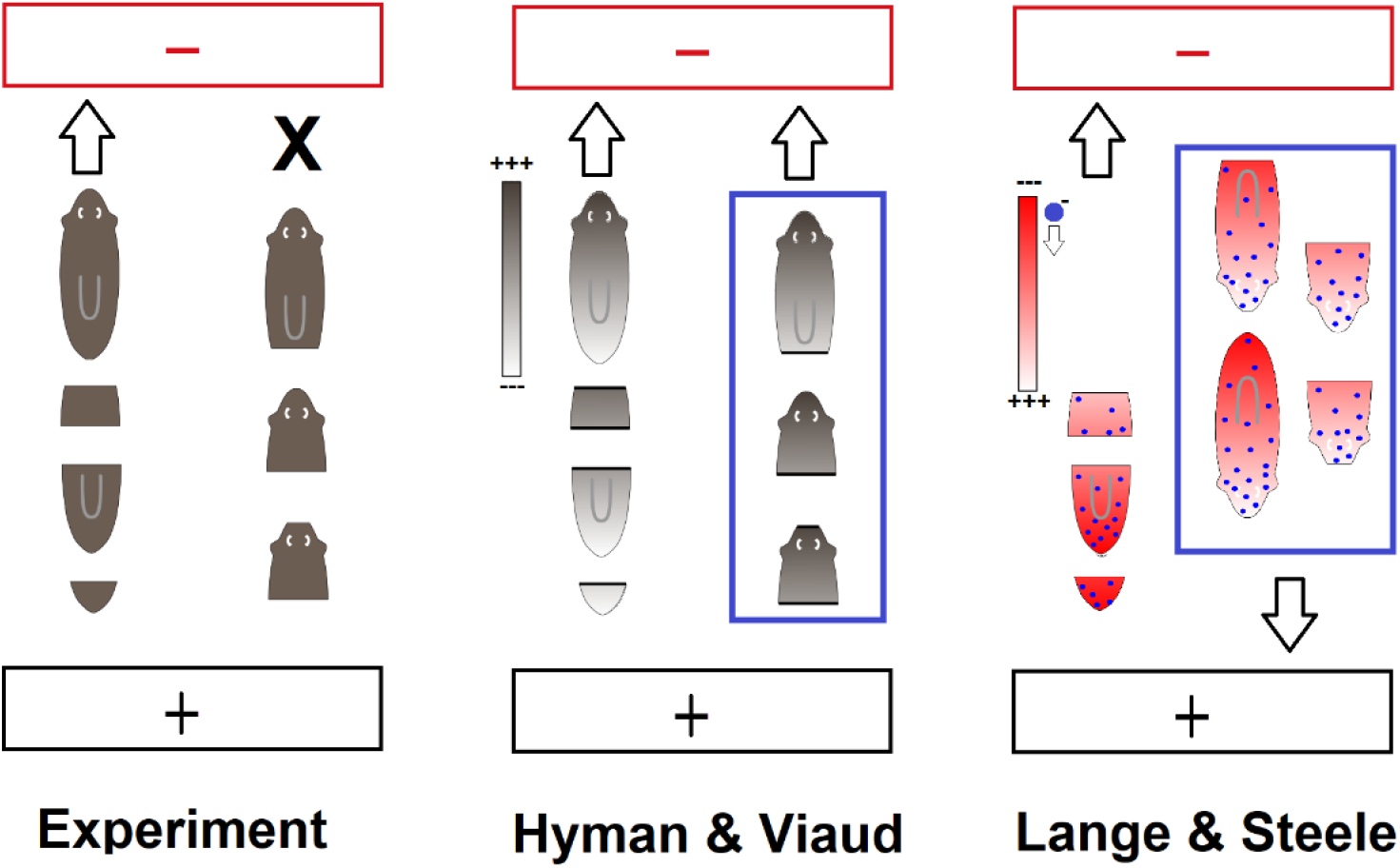
Testing proposed models against our experimental data. The cathode is indicated in red and the anode in black. Left: Results obtained in our experiments. Cathodic electrotaxis is indicated by an arrow, lack of electrotaxis is indicated by X. Middle: The Hyman & Viaud models partially explain the data but would predict cathodic electrotaxis of head fragments which was not observed in experiment. Right: The Lange & Steele model partially explain the data but would predict anodic electrotaxis of intact planarians and head fragments which was not observed in experiment. The blue boxed cases highlight which experimental data is not explained by each model.

While all models can explain the observed cathodic electrotaxis of trunk and tail fragments, the Hyman and Viaud models would predict head fragments to equally move toward the cathode, which was not observed in our experiments. The Lange & Steele model would predict intact planarians and head fragments to move toward the anode, given the presumed negative charge of the planarian head and constant production of a negatively charged morphogen in the head; however, this was also not observed in our experiments.

What distinguishes the head from the rest of the planarian body is the presence of a brain consisting of many different types of neurons organized in a bilobed neuronal network (Cebrià*et al.*, 2002; One Pagan, 2014). Viaud already proposed that differences in head and tail current sensitivity and excitation anisotropy may result from the quantity and type of neurons in each fragment (Viaud, 1952a) and proposed that the animal orients itself in the electrical field to maximize neuronal excitation. In trunk and tail fragments, the ventral nerve cords run parallel to the anterior-posterior axis, and thus could promote alignment with the external field. In contrast, in the head, neuronal connections extend in all directions (as seen from the center of the head); thus, there may not be a preferred direction of orientation and no electrotaxis is observed. One may question why we see electrotaxis in intact planarians but not in post-pharyngeally cut head pieces which are nearly of the same size. A possible explanation is the difference in head to body length ratio, which we have shown to have some effect on electrotaxis ability (**Figure 4**). We also observed that some head fragments from post-pharyngeally cut planarians show weak electrotaxis (**Figure 5**), while head fragments from pre-pharyngeal cuts never showed electrotaxis (**Figure 5**). Thus, if the body is relatively longer, the orientation of the ventral nerve cords may dominate the animal’s behavior in the electric field.

Strikingly, cathodic electrotaxis was restored in pre-pharyngeal head fragments upon decapitation. This finding that head removal enhances a behavior is in stark contrast to other stimulated planarian behaviors that require the brain. Thermotaxis, chemotaxis, and thigmotaxis are observed in intact planarians and head pieces, but not in tail pieces (Inoue, Yamashita and Agata, 2014; Inoue *et al.*, 2015). This aspect of planarian electrotaxis also differs from electrotaxis regulation in other invertebrates, where it is mediated by specific neurons in the head. The nematode *Caenorhabditis elegans* is known to move towards the cathode in response to electric fields (Shanmugam, 2017), and this behavior is disrupted when amphid sensory neurons in the head ganglia are surgically severed (Gabel *et al.*, 2007; Salam *et al.*, 2013; Chrisman *et al.*, 2016). In *Drosophila* larvae, a subset of peripheral neurons in the terminal organ at the anterior tip of the head become strongly activated when the neuronal axis becomes aligned with the direction of electric field (Riedl, 2013). In contrast, our results show that neurons in the head are not required for planarian electrotaxis; instead, their presence seems to impair electrotaxis.

Pearl’s early work in quantifying planarian behaviors and responses to stimuli (Pearl, 1903) has served as a basis for later studies linking behaviors to specific neurons or proteins, largely exclusive to the brain. The apparent divergence of electrotaxis from other stimulated behaviors that require brain control suggests that electrotaxis may be a more rudimentary behavior, like asexual reproduction or the escape response (scrunching). Since headless fragments lack other behaviors, electrotaxis may be one possible way to sense and respond to their environment. The quantitative data and methods presented here lay the foundation for future studies to dissect how headless fragments sense electric fields, its role for survival, and to determine how the head inhibits cathodic electrotaxis.

## Acknowledgments

The authors thank Veronica Bochenek for help with planarian care, Dr. William Kristan Jr. and Dr. Alex Mogilner for discussions, and Tapan Goel, Dr. Danielle Ireland, and Dr. William Kristan Jr. for comments on the manuscript.

